# Spatial arrangement of the whiskers of harbor seals (*Phoca vitulina*) compared to whisker arrangements of house mice (*Mus musculus*) and brown rats (*Rattus norvegicus*)

**DOI:** 10.1101/2024.01.15.575743

**Authors:** Matthew M. Graff, Hayley M. Belli, Sven Wieskotten, Chris S. Bresee, Yvonne Krüger, Thomas L. Janssen, Guido Dehnhardt, Mitra J.Z. Hartmann

## Abstract

Whiskers (vibrissae) are important tactile sensors for most mammals. We introduce a novel approach to quantitatively compare 3D geometry of whisker arrays across species with different whisker numbers and arrangements, focusing on harbor seals (*Phoca vitulina*), house mice (*Mus musculus*) and Norway rats (*Rattus norvegicus*). Whiskers of all three species decrease in arclength and increase in curvature from caudal to rostral. They emerge from the face with elevation angles that vary linearly with dorsoventral position, and with curvature orientations that vary diagonally as linear combinations of dorsoventral and rostrocaudal positions. In seals, this diagonal varies linearly with horizontal emergence angles, and is orthogonal to the diagonal for rats and mice. This work provides the first evidence for common elements of whisker arrangements across species in different mammalian orders. Placing the equation-based whisker array on a CAD model of a seal head enables future mechanical studies of whisker-based sensing, including wake-tracking.

**SUMMARY STATEMENT:** We quantify the three-dimensional positions and orientations of the whiskers across the face of the harbor seal, and compare this geometry with the whisker arrays of rats and mice.

## 1. Introduction

Most mammals possess specialized hairs called vibrissae, or whiskers, which extend from the mystacial pad in a grid-like array (Ahl, 1986; Muchlinski, 2010; Muchlinski et al., 2020). Different species have evolved arrays with different whisker numbers and spatial arrangements (Burns, 2009; Danforth, 1925; Grant et al., 2013; Muchlinski, 2010). To understand relationships between whisker array geometry and the whisker-based tactile signals acquired during behavior, it is important to quantify and compare whisker arrangements across species. However, existing morphometric approaches rely on landmarks or semi-landmarks (Bookstein, 2018; Shui et al., 2023), precluding comparisons across species with different whisker numbers and arrangements. New quantitative methods are required.

We recently developed a head-centered approach that uses the three-dimensional (3D) locations of whisker basepoints as parameters, enabling quantitative cross-species comparisons of whisker array morphology (Belli et al., 2018). Here we demonstrate this approach’s utility in comparing 3D whisker geometry across evolutionarily and ethologically distinct species with different whisker numbers and arrangements. We quantified the 3D geometry of the whisker array of harbor seals (*Phoca vitulina*, “seals”), which have ∼44 whiskers in seven rows, and compared the resulting equations with those for Norway rats (*Rattus norvegicus*, “rats”) and house mice (*Mus musculus*, “mice”), which have ∼30 whiskers in five rows. Seals, as aquatic predators, rely on whiskers for detecting water movements and tracking prey (Dehnhardt et al., 1998; Dehnhardt et al., 2001; Hyvarinen, 1989; Krüger et al., 2018; Miersch et al., 2011; Schulte-Pelkum et al., 2007; Wieskotten et al., 2010; Wieskotten et al., 2011). In contrast, mice and rats primarily use whiskers for direct touch on land (Brecht et al., 1997; Carvell and Simons, 1990; Welker, 1964). The head-centered approach enabled us to assess similarities and differences across species, despite different whisker numbers, and to place an equation-based model of the whisker array on a CAD model of a seal head for use in mechanical simulations and robotics experiments.

Quantitatively comparing diverse species’ arrays revealed intriguing similarities and differences. For all three species, whiskers emerge from the face with dorsoventral (“elevation”) angles that are single-variate linear with dorsoventral whisker position, and with intrinsic curvatures oriented so as to vary diagonally across the face (“twist angles”). However, these diagonals run in orthogonal directions for seals compared to rats and mice. In the seal, but not in rats and mice, whiskers emerge from the face with rostrocaudal (“horizontal”) angles that vary linearly with twist angles. We suggest that cross-species commonalities in whisker arrangements may reflect conserved mechanisms of tactile sensing, while differences could indicate ethological adaptations. Although ideally, 3D whisker array geometry would be quantified for many species, the present work provides an initial exploration into the possibility of conserved geometrical principles across different mammalian orders.

This work makes three key contributions. First, we show that harbor seal whiskers exhibit smooth, continuous gradients in arclength, curvature, and 3D orientation across the mystacial pad. We quantify these gradients and develop a corresponding CAD model of the seal head and whisker array, enabling future mechanical simulations and the development of biomechanical models for use in neuroscience and robotics. Second, we demonstrate that the head-centered approach allows, for the first time, cross-species comparisons of the 3D geometry of whisker arrays with different numbers and arrangements of whiskers. This approach establishes a foundation for quantitative comparisons of vibrissal array geometry across species. Third, we provide the first evidence for elements of a common 3D whisker arrangement across species in different mammalian orders, pointing towards conserved mechanisms of whisker-based tactile sensing, as well as evidence for cross-species differences that may reflect ethological adaptations.

## 2. Methods

### 2.1 Harbor seals and their housing and training

Three awake, behaving, adult, male harbor seals were used to quantify whisker and whisker array morphology. At the time of data collection Seal 1 (Filou, born 2006) was 5 years old, Seal 2 (Sam, born 1994) was 17 years old, and Seal 3 (Henry, born 1997) was 14 years old.

All seals resided at the Marine Science Center of Rostock University in Hohe Düne, Germany. The center is located within a marina, and consists of a moored ship (length 40 m) that serves also as an office building. The animals were kept in a mesh-surrounded 1,800 m^2^ main enclosure around the ship with a maximum depth of 6m, constantly provided with fresh seawater. Floating pontoons within the animal enclosure offer space for the animals to haul out of the water. The animals were fed approximately 80% of their daily diet (about 2 kg of freshly thawed fish cut herring or sprats per day, supplemented with vitamins) during experimental sessions as primary reinforcement.

Experiments were conducted once per day, five days per week. At the time of the experiment all animals had previously been trained to “station” from an early age, to keep their head motionless at a cued point in space for an extended duration. This established behavioral training was the only technique used to facilitate immobility during whisker measurements, therefore the tone of the facial muscles likely varied somewhat between individuals and between scans. Although we have no measurement of degree of activation of vibrissal-related facial muscles, we qualitatively judged that vibrissae were in a neutrally relaxed position (neither retracted nor protracted) and generally in equivalent positions across scans and across animals, based on visual assessment of the data. Quantification of whisker emergence angles (*Results*) later confirmed the similarity across individuals. The reproducibility in emergence angles is unsurprising due to the highly stereotyped nature of the well-trained and frequently repeated stationing behavior.

Animals were not required to station for a fixed duration. Instead, stationing was completely voluntary and an animal could choose to return to the enclosure at any time. Typical stationing times were approximately 30 – 40 sec, permitting at least one complete 3D scan, or several 2D photographs, to be obtained during the stationing event.

All experiments performed at the Marine Science Center of Rostock University were conducted in accordance with the European Communities Council Directive of 24 November 1986 (86/609/EEC), the directive 2010/63/EU and the Directive of the §7 and §11 TierSchG of the German Protection of Animals Act (Tierschutzgesetz).

### 2.2 Data acquisition and extraction

#### 2.2.1. 2D data acquisition: photographs of individual whiskers

We took 127 two-dimensional (2D) photographs of 52 unique whiskers. All photographs were taken when animals voluntarily and transiently chose to emerge from the sea enclosure around the ship, so whiskers were in a hydrated state. Hydration state is important when estimating intrinsic whisker curvature because informal observation indicates that a whisker’s shape changes substantially as it dehydrates, on the time scale of ∼20 minutes.

Because whiskers could not be plucked from the animal, photographs of the whiskers were taken against a black, hard-plastic background. A camera-frame system (Fig. 1A) was constructed to ensure that the whiskers lay flat on the background with no distortion of their natural shape. The frame system, constructed of 80/20^®^, also ensured a fixed distance between the camera (Panasonic, Lumix DMC-FZ20) and the background, and resulting photographs had a resolution of ∼200 microns/pixel. Length scales (thin plastic rulers) were affixed rigidly to the background.

**Fig. 1:**
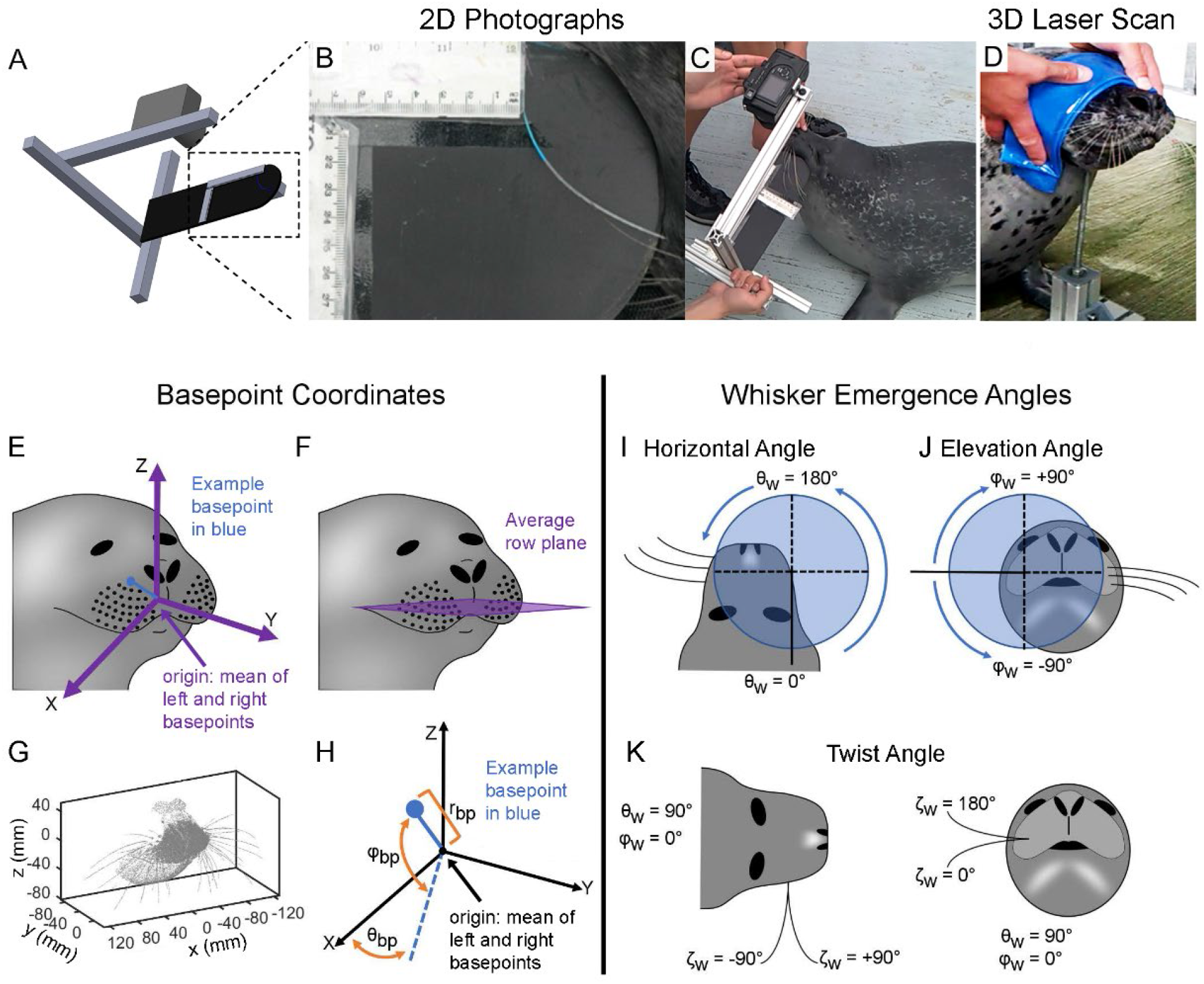
After imaging the head and vibrissal array of a harbor seal in 3D and 2D, coordinate systems for whisker basepoints and emergence angles were chosen. **(A)** A schematic of the custom-built camera frame system used to obtain 2D photographs of the whiskers. The orientation of the background was manually adjusted so that the whisker of interest lay flat on the background. **(B)** An expanded view of the camera frame system shows an exemplary 2D photograph of a whisker identified with non-toxic watercolor paint. **(C)** A photograph shows how the custom-built camera frame system was used in practice. **(D)** A blindfolded seal is shown perching its head on a chinrest to permit 3D laser scanning. **(E)** Schematic of standard position and orientation of the vibrissal array. The origin (0,0,0) is the mean position of the basepoints of left and right arrays. The right basepoints surround the positive x-axis. The blue line and dot indicate one example of a basepoint location, for comparison with panel H. **(F)** The horizontal (x-y) plane is defined by the average row plane of the basepoints. **(G)** An example point cloud of a seal’s face in standard position and orientation. **(H)** Each basepoint location is described in spherical coordinates (θbp, ϕ_bp_, r_bp_). The angle θ_bp_ ranges from -90° (caudal) to +90° (rostral), where θ_bp_ = 0° lies in the x-z plane. The angle ϕ_bp_ ranges from -90° (ventral) to +90° (dorsal), where ϕ_bp_ = 0° lies in the x-y plane. The basepoint coordinates of one example whisker (blue dot) are shown. **(I)** The variable θ_w_ describes the rostrocaudal angle at which the proximal, approximately-linear portion of the whisker emerges from the mystacial pad. Values range from 0° to 360°. **(J)** The variable ϕ_w_ describes the dorsoventral angle at which the proximal, approximately-linear portion of the whisker emerges from the mystacial pad. Values range from -90° to 90°. **(K)** The variable ζ_w_ describes the whisker’s twist about its own axis. Values range between ±180°. Photos in (B-D) have been brightened and contrast enhanced.

The 2D photographic procedure was as follows. A select number of whiskers were marked using non-toxic watercolor paint. The background was then manually held against the face of the seal in an orientation such that one selected whisker lay flat against the background without any shape distortion. Although several whiskers (painted different colors) were often visible in each photo, the frame was oriented so that one specific whisker lay completely flat. During analysis, we could be confident that we captured the entire arclength of the whisker because the tip of each whisker was clearly visible with the naked eye against the dark black background in the digital photograph. An example of a typical 2D photo of a whisker with a blue stripe is shown in Fig. 1B. The photo in Fig. 1C shows how the frame was oriented by hand to position the whisker onto the background. A log of the correspondence between painted color and whisker identity was recorded as the photographs were taken, and the images of the seal whiskers were then imported into Matlab™ (The Mathworks, Natick, MA) for measurement.

#### 2.2.2. 2D data extraction: whisker arclength and intrinsic curvature

The 2D photographs allowed us to quantify two parameters: whisker arclength (S) and intrinsic curvature (A).

Whisker arclength refers to the length measured along the curve of the whisker, distinct from the “base-to-tip” length, which is the chord length (straight-line distance)between the whisker’s base and tip. To quantify arclength, whisker shapes were manually traced by clicking along the approximate centerline of each whisker, from base to tip. The traced points were then up-sampled via linear interpolation to 100 points and smoothed with a moving average filter using a 15-sample window. All traces were overlaid onto the original photographs and visually inspected for accuracy. The arclength of each whisker was calculated by summing the lengths of the segments composing the traced image. Finally, all traces were converted to real-world units using the scale bars attached to the background (Fig. 1B).

To quantify intrinsic whisker curvature, we aimed for the simplest and most intuitive description possible. Following established methods (Belli et al., 2018; Bresee et al., 2023; Knutsen et al., 2008; Towal et al., 2011), each whisker was characterized by a single “curvature coefficient,” based on a polynomial fit. Each seal whisker was fitted with both quadratic and cubic curves, then visually assessed for goodness of fit. Cubic fits were superior to quadratic fits, as they accommodated the near-linear proximal portion while also capturing most of the distal curvature. In its most general form, a cubic polynomial has four coefficients: f(x) = Ax^3^ + Bx^2^ + Cx + D. For simplicity, we rotated each whisker to ensure that the B, C, and D coefficients were zero, and performed a least-squares regression to the curve f(x) = Ax^3^. This alignment procedure requires the proximal portion of the whisker to be aligned with the x-axis. To achieve this alignment, the whisker was divided into 100 nodes, with the base point placed at the origin (0,0), and the next proximal node (∼1% of the whisker arclength) aligned with the x-axis. This method has the advantage of requiring only a single coefficient, with an intuitive meaning, to describe the intrinsic curvature of the whisker.

Previous studies (Dougill et al., 2023; Luo and Hartmann, 2023; Towal et al., 2011) have employed more complex and detailed approaches to quantifying whiskers’ intrinsic curvatures. These studies suggest that a fractional exponent model (Luo and Hartmann, 2023) or Cesaro fits (Dougill et al., 2023; Towal et al., 2011) may provide more precise estimates of whisker shape.

Because the present study was performed in awake behaving animals, the resolution of the 2D images did not permit analysis of whisker diameter, elliptical cross-sections, taper, or the 3D angles that quantify undulations along the whisker length. Future work should systematically quantify these parameters.

#### 2.2.3. 3D data acquisition: laser scans

The head and vibrissal array of each seal were scanned using a (3D) volumetric scanner, resulting in 361 3D whisker images containing 171 unique whiskers. A Konica Minolta Vivid 900 non-contact, 3D laser scanner was used for these scans. Scan resolution depends on distance to the subject and scan duration, parameters that necessarily varied because we were working with awake behaving animals. The scanner can acquire up to ∼300,000 points in 2.5 seconds within a ∼10×10 cm^2^ box at a range of ∼5 cm, or within a ∼100×100 cm^2^ box at a range of ∼75 cm. Each seal was trained to pose its head on a semicircular stand (a chinrest) for less than one minute, allowing for one to two scans per session. During scanning, the seal’s eyes were covered with a thick, opaque rubber mask. An example of a blindfolded seal resting its head on the chinrest is shown in Fig. 1D. The software package Rapidform™ XOR2 was used to control the scanner from a laptop computer. The scanner output was a digitized point cloud which included portions of the seal’s head, neck, and whiskers.

#### 2.2.4. 3D data extraction: alignment of multiple laser scans

Because the seal’s head and whisker arrays were larger than the 3D scanner’s field of view, multiple scans of the same animal had to be aligned and merged in order to construct a 3D representation of the whisker array.

We first performed a coarse alignment of the point clouds within Rapidform™. Data points associated with the head or neck, or with scanner noise, were manually removed. The most complete point cloud for each seal was chosen as the reference scan. All other scans were rotated and translated to align to the reference scan using facial landmarks, which included whisker basepoints, the nostrils, and face and mouth creases. Whiskers were identified by their location within the array (Dehnhardt and Kaminski, 1995) as well as by comparing the point clouds to photographs of the full mystacial pad of the seal. All data points corresponding to each individual whisker were saved into their own point clouds.

Next, continuing to work within Rapidform™, the whisker basepoint was identified for each whisker point cloud. The basepoint was determined by finding the intersection between the whisker and the skin. In nearly all cases this whisker-skin intersection was clearly visible and the basepoint was determined as the centroid of 5 to 15 points at the intersection. Scans in which the basepoint could not be clearly identified were excluded from further analysis.

At this stage of the alignment process, each 3D scan of a particular seal now consists only of point clouds of identified whiskers, and each whisker point cloud has an average basepoint. All non-whisker related points have been removed, ensuring that the automated alignment described in the next step included only the appropriate whisker-related points.

Finally, a fine alignment of these “whisker-and-basepoint-only” scans. As before, the scan with the largest number of basepoints was used as the reference scan. An optimization was performed using the “fminsearch” function in Matlab™ in which each scan was rotated and translated into alignment with the reference scan. The objective function for the minimization consisted of the summed distances of all matching basepoints between the reference scan and each individual scan. This final step produced well-aligned 3D scans form a single seal. Scans from multiple individuals were never aligned.

### 2.3. Choose origin and horizontal plane

After data acquisition and extraction, the next step was to choose an origin and horizontal plane based on selected features of the animal’s face and/or skull (Figs 1E – 1H). Because the present work focused on the vibrissae, we chose the mean of all whisker basepoints as the origin, and the “average row plane” (Towal et al., 2011) to define horizontal.

To place the head into a standard position, the aligned point clouds were translated so that the origin was located at the mean position of all right and left whisker basepoints. This puts the origin inside the animal’s head, near the center of the muzzle (Fig. 1E). This procedure was implemented using “matched” basepoints between right and left sides; in the case that a basepoint was missing on one side, its non-missing complement was omitted from the calculation of the origin. For seals with multiple measurements of the same basepoint, the mean of the points was used for the alignment procedures.

To put the head into a standard orientation, we used the “average row plane” of the whiskers to define horizontal (Fig. 1F). We performed a roll-yaw-pitch (y-z-x-axis) rotation sequence. The roll rotation (about the y-axis) and the yaw rotation (about the z-axis) placed the center of the right array on the positive x-axis and the center of the left array on the negative x-axis. The “center” of the right (left) array is defined as the mean of all basepoint positions on the right (left) side of the animal.

The pitch rotation (about the x-axis) aligned the average whisker row plane with the x-y (horizontal) plane. To find the average whisker row plane, individual planes were fitted to the basepoints of the combined left and right whisker rows using least squares and producing six normal vectors – one for each of the six whisker rows B - G. The A row of whiskers was excluded because it contained an insufficient number of points to define a plane. The maximum sum-squared-error between the basepoints and its fitted plane was 90.2 mm^2^ (maximum residual was 5.5 mm). The mean of the unit normal vectors describes the average row plane (Fig. 1F). The animal’s head was then pitched about the x-axis so that the average row plane was parallel with the x-y plane. The final position of the seal point cloud is shown in Fig. 1G.

### 2.4. Quantify whisker basepoint locations and the surface area of the array

The seal whisker array is organized in a grid-like pattern that includes 7 rows and 9 columns (Dehnhardt and Kaminski, 1995). In contrast, mice and rat whisker arrays are organized into 5 rows and 7 columns (Woolsey and Van der Loos, 1970). .In order to permit comparisons across animals with different numbers and arrangements of whiskers, we quantified the morphology of the seal whisker array in terms of the coordinates of the whisker basepoints (*r*_*bp*_, *θ*_*bp*_, ϕ_*bp*_), rather than row and column identity.

Basepoint coordinates were determined within a modified spherical coordinate system (Fig.1H) with the origin located at the mean position of all the basepoint locations. Spherical coordinates are used because they adapt most naturally to the (roughly spherical) geometry of an animal’s head. For each whisker, the variable θ_bp_ describes the rostro-caudal location of the basepoint with respect to the positive x-axis. A positive value of θ_bp_ indicates that the whisker is more rostral, while a negative angle indicates that the whisker is more caudal. The variable ϕ_bp_ describes the dorsoventral location of the basepoints with respect to the x-y plane. A positive value of ϕ_bp_ indicates a more dorsal basepoint location, while a negative coordinate indicates a more ventral basepoint location. The third basepoint coordinate is the radius (r_bp_), i.e., the straight-line distance between the origin and the basepoint. Scaling the basepoint radii scales the size of the seal’s face at each whisker location.

To generate an estimate of surface area of the whisker array we first found the mean position of each whisker basepoint across all seals. We then found the residuals of each basepoint (from individual seals) from its corresponding mean. Next, we manually connected three adjacent basepoint identities to form a trimesh of non-overlapping triangles that covered the entire whisker pad surface. The whisker array area was computed as the sum of the areas of these triangles. The range for the area was computed by computing the surface area using the mean plus and minus the extreme values of the residuals in all directions. The same procedure was performed for rats and mice, using data from Belli et al., 2016 and Bresee et al., 2023.

### 2.5. Quantify each whisker’s orientation on the face with three emergence angles

The orientation of each whisker on the face is given by the three angles at which it at which it emerges from the mystacial pad (Towal et al., 2011). These angles, illustrated in Figs. 1I – 1K, are termed the “emergence angles,” and in the present work, we describe them using Tait-Bryan angles applicable to the right array.

The horizontal angle θ_w_ describes the rostrocaudal orientation of the proximal portion of the whisker. A θ_w_ angle of 180° is directed rostrally while an angle of 0° is directed caudally (Fig. 1I). The elevation angle ϕ_w_ describes the dorsoventral orientation of the proximal portion of the whisker. A ϕ_w_ angle of 90° points the whisker dorsally while an angle of -90° points it ventrally (Fig. 1J). Whiskers have an intrinsic curvature and therefore they require a third angle, ζ_w_, to describe the roll about their own axis. For a whisker with θ_w_ = 90° and ϕ_w_ = 0°, a ζ_w_ angle of 90° will curve concave forward and -90° concave backward (Fig. 1K). In contrast, a ζ_w_ angle of 180° will curve concave upward, and 0° concave downward (Fig. 1K). Note that the directionality of the whisker’s concavity in the head-centered coordinate system is defined by all three orientation angles, not just ζ_w_.

We next had to determine each whisker’s emergence angles based on the data in its 3D point cloud. To find the emergence angles, an optimization was performed using the built-in Matlab™ function “fmincon.” The parameters for the optimization were θ_w_, ϕ_w_, ζ_w_, along with S and A. Constraints on the angles of emergence in the optimization were: θ_w_ = [0, 360], ϕ_w_ = [-90, 90], ζ_w_ = [-180, 180], S = [straight distance from base to tip, lower bound + 50mm], curvature coefficient A = [0 to 1]. The lower bound for S was chosen to be the linear distance between the base and tip (the smallest S could ever possibly be to connect those two points) and 50 mm longer than the lower bound. Beginning with the whisker point cloud in standard position and orientation, the individual points within the cloud were sorted by distance from the basepoint.

The x, y, z coordinates of all the points were then smoothed with a moving average filter using a 5-sample window. The whisker was resampled into 500 micron segments and translated to have the base at the origin. The routine subsequently varied five variables, θ_w_, ϕ_w_, ζ_w_, S, and A to fit an idealized whisker model to the whisker point cloud. The idealized model started with the base at the origin (θ_bp_ = 0, ϕ_bp_ = 0, r_bp_ = 0), and the proximal section of the whisker was coaxial to the negative y-axis (θ_w_ = 0, ϕ_w_ = 0). The model’s orientation was concave down (ζ_w_ = 0). To align the idealized model with the whisker point cloud, an Euler rotation sequence was performed. The order of the rotations was y-axis (roll, ζ_w_), x-axis (pitch, ϕ_w_), z-axis (yaw, θ_w_). All rotations were extrinsic around the global y-x-z axes. Additionally, the arc-length and curvature were optimized to best match the idealized model to the point cloud. The optimization’s objective function minimized the mean sum squared distance between the whisker point cloud and the points in the idealized whisker model. The values of θ_w_, ϕ_w_, ζ_w_ found for the best match were taken to be that whisker’s angles of emergence.

It is important to note that the emergence angles were measured from 3D scans obtained with the whiskers in a neutrally relaxed position. Given that the seal whisker array is mobile, these angles reflect only one possible position for the whiskers. Future work could use the 3D model developed here to simulate and quantify the effects of whisker motion and angle changes on the mechanics of sensory acquisition behaviors and wake following. In the *Discussion*, we assess only the neutral, relaxed emergence angles presented in *Results*.

### 2.6. Use whisker basepoint locations as independent variables to develop equations for whisker and array geometry

We aimed to develop equations that relate whisker arclength (S), intrinsic curvature (A), the radial basepoint (r_bp_), and the angles of emergence (θ_w_, ϕ_w_, ζ_w_) to the locations of the whisker basepoints θ_bp_ and ϕ_bp_. Throughout the work, all equations and figures are presented in terms of whisker arrays on the animal’s right side. To obtain results for the left side, all equations just be mirrored across the y-z plane.

#### 2.6.1. Equations for 2D geometry: Arclength and intrinsic curvature as functions of θ_bp_ and ϕ_bp_

The 3D laser scans yielded a total of 171 unique images of whiskers from three seals. However, when these 171 whiskers were compared across individuals, several whiskers were found to have unusually short arclengths given their identity. The short arclength was attributable to two causes. First, in some scans, the 3D scanner did not have sufficient spatial resolution to capture the full length of the whisker. Second, the scans were performed during the summer months, when two seals were undergoing vibrissal renewal and replacement (Hirons et al., 2001) and some of the whiskers had not fully regrown.

Rather than fit a model that under-predicted S and inaccurately predicted A, we excluded the data for S and A from the 3D scans, and constructed multivariate linear regressions models for S and A as functions of θ_bp_ and ϕ_bp_ using only the 2D dataset of 52 unique whiskers. For these 2D data, scan quality did not contribute significantly to measurement error in the arclength, as was the case for the 3D scans. The unusually short 2D measurements were attributable only to vibrissal renewal and replacement. To determine outliers due to vibrissal renewal and replacement, any whisker from the 2D data (from a particular seal) that was more than 30 mm shorter than the longest whisker (across all seals, for that whisker identity) was excluded. This threshold captures values greater than two standard deviations from the mean whisker length. Eight of the 52 2D traces were excluded based on this criterion, and the remaining 44 2D whiskers were then used to find relationships between S, A and θ_bp_ and ϕ_bp_.

Among the 44 whiskers used in constructing the 2D model, 27 came from Seal 1, 13 from Seal 2, and four from Seal 3. Using the reduced set of 44 whiskers, first-order models for the log-transformed 2D arclength (S) and cubic coefficient (A) were fit as functions of θ_bp_ and ϕ_bp_. Because 2D traces did not provide basepoint coordinate measurements, the average basepoint coordinates from the 3D scans were matched to the whisker identity of each 2D trace.

Even though there are fewer 2D measurements than 3D measurements for S and A, we can be confident that they capture the full length of the whisker and will therefore more accurately depict the cubic coefficient. Therefore, results for arclength and cubic coefficient (Fig.2D-I and Eqns 2-6) are based only on the 2D data.

#### 2.6.2. Equations for 3D geometry: Radial basepoint (r_bp_) and angles of emergence (θ_w_, ϕ_w_, and ζ_w_) as functions of θbp and ϕbp

As described for the analysis of 2D geometry, some whiskers from the 3D images were found to have unusually short arclengths. However, we did not remove any whiskers when finding equations for r_bp_ and θ_w_, ϕ_w_, ζ_w_ as functions of θ_bp_ and ϕ_bp_, because the basepoint will not change, and because we fully expect whiskers at various stages in the growth cycle (i.e., with various lengths) to retain the same emergence angles.

We aimed to construct multivariable linear regression models for r_bp_, θ_w_, ϕ_w_, and ζ_w_ as functions of θ_bp_ and ϕ_bp_. Rather than fit a model directly to the entire dataset of 171 whiskers and potentially overfit the data, we wanted to first find the lowest order model that could fit all three seals individually, and then apply a model of same order to the entire dataset of 171 whiskers. We therefore constructed models using the well-established method of forward selection and evaluation based on the Akaike information criterion (AIC), first finding the best fit model for each individual and then applying the most common model form to all individuals together.

We first divided the data from 171 whiskers for r_bp_, θ_w_, ϕ_w_, and ζ_w_ into three subgroups by seal identity. Creating individual models for each seal ensured that we did not overfit by treating each whisker from an individual array as an independent measurement. For each of the three seals, we made histograms for r_bp_, θ_w_, ϕ_w_, and ζ_w_. All 12 histograms were found to be normally distributed. Linear regression models were constructed for each seal to test whether r_bp_, θ_w_, ϕ_w_, and ζ_w_ were univariately associated with θ_bp_ and/or ϕ_bp_. We tested the hypothesis that the θ_bp_ and/or ϕ_bp_ regression coefficients were equal to zero. If the p-value for each independent variable coefficient was less than 0.05 we tested a second order model (square of θ_bp_ and/or square of ϕ_bp_), but only accepted the higher order model when it also had a p-value <0.05 and an AIC of at least 2 points lower than the lower order model. If the second order model was significant and supported by a lower AIC, a third order model (cube of θ_bp_ and/or cube of ϕ_bp_) was tested, and accepted if it had a p value <0.05 and at least a 2-point lower AIC than the quadratic model. We did not find any models for whiskers from an individual seal to have independent variable predictors greater than third-order.

Finally, after finding the best fitting equations for each individual, the lowest order model among the three seals with significant regression coefficients was selected as the final model. These steps allow us to write equations for *S, A, θ*_w_, ϕ_w_, *ζ*_w_ in terms of θ_bp_ and ϕ_bp_, so that the equations describing the whiskers and whisker array geometry can be compared across different species.

We note that this same approach could in principle be used to parameterize the locations of facial and skull features, as well as the major and minor diameter axes and undulation angles.

### 2.6 Creating a CAD model of the seal head

To create a CAD model of the seal’s head we used 3D laser scans from Seal 2 that included the muzzle, face, and neck. The scans were taken at four different orientations: one orientation focused on the nose and top of the head, a second focused on the lower jaw and neck, and the third and fourth on the right and left muzzles, respectively. Visualization of the scans revealed that Seal 2 had its mouth open by a different amount during the scans of the right and left muzzles. Because the mouth position changed between scans, the lower jaw could not be accurately aligned with both right and left muzzles simultaneously: the jaw could be aligned either with the right muzzle or the left, but not both. To solve this problem, the left muzzle was mirrored to the right, allowing jaw-muzzle alignment without any manual editing. After mirroring the left muzzle, all scans were meshed, aligned, and merged within the software Geomagic™. This procedure yielded a single surface mesh that included the seal’s neck, lower and upper jaws, nostrils and snout, portions of the top of the seal’s head, and mirror-symmetric muzzles.

However, because the live seals were blindfolded during the 3D laser scans, the mesh from Seal 2 contained only limited information about the locations of the eyes. We therefore performed a 3D laser scan of the head of a deceased adult, male harbor seal obtained from *Allied Whale* at the College of the Atlantic in Bar Harbor, Maine. The tissue transfer was approved in advance by the Institutional Animal Care and Use Committee of Northwestern University and the work was performed with authorization from the U.S. National Oceanic and Atmospheric Administration’s (NOAA’s) National Marine Fisheries Service (NMFS) in accordance with the U.S. Marine Mammal Protection Act regulations. A Keyence Romer arm 7525 S1 was used to take multiple scans of the head, which were then merged and meshed within Geomagic™.

The mesh of Seal 2 was aligned with the mesh of the deceased seal, using the noses, muzzles, and the nasal ridge as landmarks. The alignment required scaling the mesh of the deceased seal by a factor 0.98 about its center of mass. After alignment, all parts of the mesh from the deceased seal were removed except for the eyes and portions of the top of the head. Finally, to ensure a closed volume and a solid CAD model, the merged surface was smoothed behind the muzzles up towards the eyes. To summarize, the final CAD model was obtained entirely from scans of a live seal (Seal 2), with the exception of the eyes, part of the top of the head, and small, smoothed regions behind the two mirror-symmetric muzzles.

### 2.7. Comparison of the 3D geometry of the harbor seal whisker array with the 3D geometry of house mice and brown rats

To compare the 3D geometry of the harbor seal whisker array with the geometries of the whisker arrays of rats and mice, we used data from two previously published studies. Rats and mice both have ∼27 large macrovibrissae on both sides of their face, arranged into five rows (typically given the letters A – E) and six or seven columns, depending on the row (typically numbered 1 – 7). The four most caudal whiskers (designated by the letters *α* through *δ*) have basepoints that lie between the five rows (Woolsey and Van der Loos, 1970).

Data for the rat were obtained from Belli et al., 2018, which developed equations for the whisker array based on a total of 518 macrovibrissae from nine female Long Evans rats between the ages of 5 and 36 months. 3D geometries were obtained either from a 3D laser scan or with a Microscribe*™*, a passively movable mechanical arm with five degrees of freedom that can be used to “trace” along the whisker length. The user moves a probe tip at the end of the arm to a chosen location and that location is recorded as (*x, y, z*) coordinates relative to a pre-defined origin. Data for the mouse were obtained from Bresee et al., 2023. A total of 467 macrovibrissae from eight mice were collected for the present work. All animals were male C57BL/6J mice between the ages of 6 and 8 weeks.

To generate an estimate of surface area of the whisker array we first found the mean position of each whisker basepoint across all seals. We then found the residuals of each basepoint (from individual seals) from its corresponding mean. Next, we manually connected three adjacent basepoint identities to form a trimesh of non-overlapping triangles that covered the entire whisker pad surface. The whisker array area was computed as the sum of the areas of these triangles. The range for the area was computed by computing the surface area using the mean plus and minus the extreme values of the residuals in all directions.

As noted previously (Bresee et al., 2023), a limitation of the comparison between mice and rats is that Bresee et al. (2023) used exclusively adult male mice, while Belli et al., 2018 used only adult female rats. However, we expect any potential differences between females and males of the same species and similar size to be smaller than the spatial resolution of the scanning approaches.

As stated in Belli et al., 2018, all procedures for rats were approved in advance by the Animal Care and Use Committee of Northwestern University. As stated in Bresee et al., 2023, all procedures for mice were approved in advance by the Animal Care and Use Committee of Northwestern University.

## 3. Results

### 3.1. Basepoint coordinates

Dehnhardt and Kaminski (1995) characterized the qualitative shape of the seal whisker array and identified whiskers within it (Fig. 2A). Plotting ϕ_bp_ vs. θ_bp_ yields a quantitative depiction of the array that closely resembles the qualitative description (Fig. 2B). The discontinuity in row direction between columns 4 and 5 is maintained, although evidenced as a gradual bend rather than a sudden break.

**Fig. 2:**
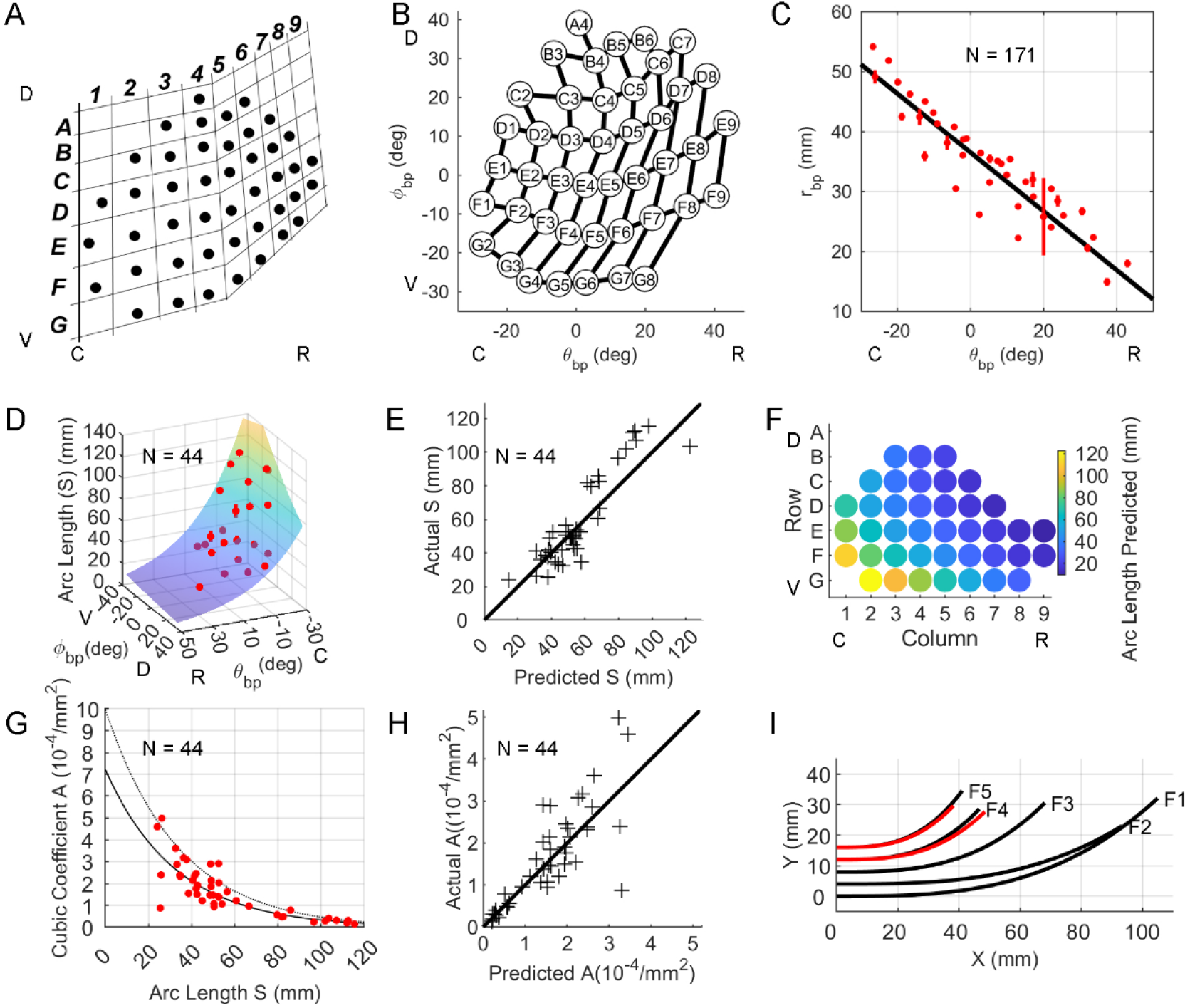
Locations of whisker basepoints and their relationship to 2D whisker geometry. **(A)** Qualitative description of the shape of the harbor seal whisker array. Figure recreated from (Dehnhardt and Kaminski, 1995). **(B)** Plotting ϕ_bp_ against θ_bp_ quantifies array geometry in a manner that closely resembles the qualitative description in (A). **(C)** Radial basepoint coordinate (r_bp_) is plotted as a function of θ_bp_ for N = 171 basepoints. Black line represents Eqn 1. Mean values of r_bp_ when grouped by Row and Column identity are shown as red dots; red vertical bars indicate standard errors. **(D)** Arclength (S) is plotted as a function of θ_bp_ and ϕ_bp_. The surface represents Eqn 2. Mean values of S when grouped by Row and Column identity are shown as red dots; red vertical bars indicate standard errors. **(E)** Experimentally measured values for arclength (S) are compared with values predicted from Eqn 2. Black diagonal line indicates equality. **(F)** A colormap illustrates variations in arclength predicted by Eqn 2 across the array. **(G)** Cubic curvature coefficient (*A)* can be described as an exponentially decreasing function of arclength. Solid black line represents Eqn 4 and dashed gray line represents Eqn 5. Red dots are experimentally measured values of A. **(H)** Experimentally measured values for curvature coefficient (A) are compared with values predicted from Eqn 4. Black diagonal line indicates equality. **(I)** Black traces show shapes of the F-row whiskers from a single seal. Whiskers F4 and F5 from two other seals are shown in red to illustrate shape variability. Abbreviations: caudal (C); rostral (R); ventral (V); dorsal (D). Subplots D, E, G, and H contain data from 44 whiskers.

The basepoint coordinate θ_bp_ was used as an independent variable to quantify the radial coordinate of the basepoints (r_bp_). Results for individual seals, with confidence intervals, are shown in Supplementary Table S1 and indicate that r_bp_ depends only on θ_bp_:

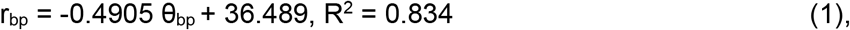

where θ_bp_ is in degrees and r_bp_ is in millimeters.

Eqn 1 indicates that r_bp_ decreases from caudal to rostral, a trend depicted in Fig. 2C. The radial basepoint’s independence from ϕ_bp_ aligns with the observation that the seal’s mystacial pad is nearly vertical along the dorsoventral axis.

### 3.2. 2D shape: arclength and curvature

Basepoint coordinates θ_bp_ and ϕ_bp_ were used as independent variables to quantify variations in arclength (*S*) and intrinsic whisker curvature (*A*) across the array. When averaged across all animals, S was best described as an exponential function of θ_bp_ and ϕ_bp_:

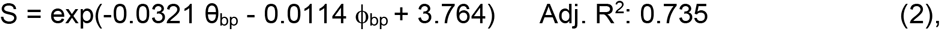

where θ_bp_ and ϕ_bp_ are in degrees and S is in millimeters.

The exponential decrease of S with θ_bp_ and ϕ_bp_ is shown in Fig. 2D. The surface represents Eqn 2 and red points are the mean value of S when grouped by whisker identity +/- standard error. Fig. 2E compares the arclength values predicted by Eqn 2 with experimental measurements. The model underestimates the length of whiskers longer than ∼60 mm, and whiskers shorter than ∼60 mm are broadly clustered about the diagonal, suggesting that the arclength prediction might improve with more data. Finally, Fig. 2F shows predicted values for S across the whisker array.

Whisker intrinsic curvature was quantified using the same 44 whiskers used to quantify arclength. The shapes of the seal’s whiskers were well described by the function y = Ax^3^, with the coefficient A termed the “cubic curvature coefficient.” When quantified in terms of basepoint parameters the value of A was not significant in ϕ_bp_, so Eqn 3 expresses A only as a function of θ_bp_:

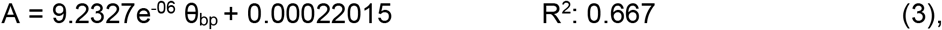

where θ_bp_ has units of degrees and A has units of 1/mm^2^.

However, further investigation found that whisker curvature was significantly better described as an exponential function of arclength rather than as a linear function of θ_bp_:

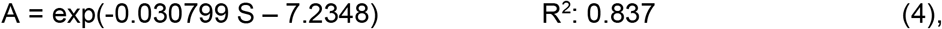

where S has units of millimeters and A has units of 1/mm^2^. The 95% confidence intervals for the arclength coefficient and intercept are [-0.0350, -0.0266] and [-7.4968, -6.9727], respectively. Eqn 4 indicates that A will typically be larger for shorter whiskers than for longer whiskers.

Fig. 2G shows Eqn 4 as a solid black line plotted over the experimental data in red. Although the fit is excellent for whiskers with arclengths greater than ∼55 mm, there is considerable scatter for shorter whiskers. This increased scatter is easily observed when experimental values for A are compared with those predicted by Eqn 4 (Fig. 2H). These results closely resemble those found for rats and mice, whose rostral whiskers also exhibit high variability in intrinsic curvature (Belli et al., 2018; Bresee et al., 2023; Towal et al., 2011). Following the approach used for rodents, we found the upper bound on the relationship between A and S. This bound is shown as a thin, dashed, gray line in Fig. 2G, and follows the equation:

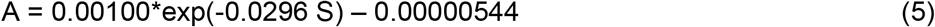

The upper bound limits the y-coordinate of the whisker’s tip, perhaps limiting total whisker “height.” (Belli et al., 2018). Finally, to provide visual intuition for S and A, traces of the F-row of whiskers from a single seal (Seal 1) are shown in black in Fig. 2H. A sense for the magnitude of the variability is provided in the red traces of Fig. 2H, which show whiskers F4 and F5 obtained from the two other seals.

### 3.3. 3D shape: angles of emergence

Basepoint coordinates θ_bp_ and ϕ_bp_ were used as independent variables to quantify the angles of emergence of the whiskers across the array. Best fit equations for θ_w_ and ζ_w_ were linear in both θ_bp_ and ϕ_bp_, while the best fit equation for ϕ_w_ depended only on ϕ_bp_:

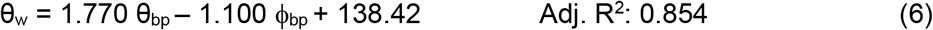

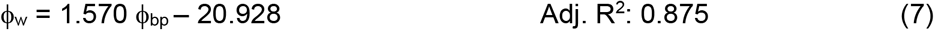

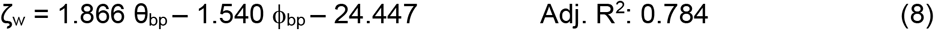

In Eqns 6, 7, and 8, all variables have units of degrees. Lines 2 - 5 of Supplementary Table 1 provide data for individual seals, with confidence intervals.

Fig. 3ABC compares these equations with experimentally-measured values. The horizontal emergence angle (θ_w_) increases from caudal to rostral across the array and from dorsal to ventral (Fig. 3A). The elevation angle ϕ_w_ increases from ventral to dorsal, with no significant change in the caudal-rostral direction (Fig. 3B). Notably, the ϕ_w_ angles are not symmetric about 0 deg, indicating that the whiskers are skewed to point downward. The twist angle, ζ_w_, increases from caudal to rostral across the array and from dorsal to ventral (Fig. 3C).

**Fig. 3:**
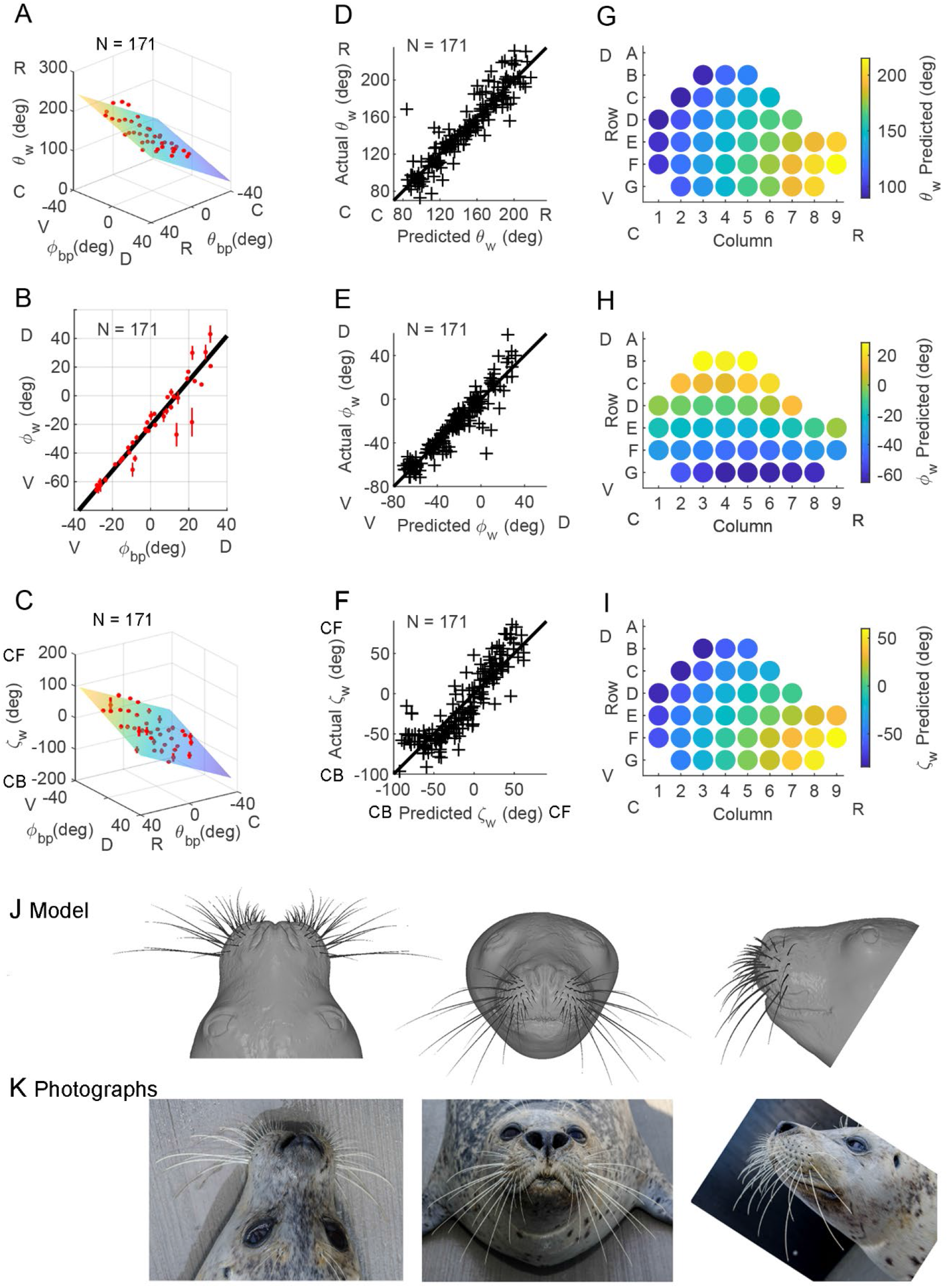
Quantifying the relationship between whisker angles of emergence and basepoint coordinates allows construction of a CAD model of the whisker array. **(ABC)** The subplots show the horizontal angle of emergence (θ_w_) fit to Eqn 6, the elevation angle of emergence (ϕ_w_) fit to Eqn 7, and the twist angle of emergence (ζ_w_) fit to Eqn 8. For all plots, mean ± standard error when grouped by whisker identity is shown in red. **(DEF)** The subplots show actual values of θ_w_, ϕ_bp_, and ζ_w_ compared with values predicted from Eqns 6, 7, and 8. All subplots reveal relatively uniform dispersion of actual vs. predicted values about y = x, indicating that correct models were selected. **(GHI)** The subplots show how values for θ_w_, ϕ_bp_, and ζ_w_ will vary across the array, as predicted by Eqns 6, 7, and 8. Panels A - E show data for N = 44 seal whiskers photographed in 2D. Panels G-O show data for N = 171 seal whiskers scanned in 3D. **(J)** Eqns 1 – 8 can be used to develop a CAD model of the seal whisker array, shown superposed on a model of the seal’s head. The head model was created using 3D laser scans of Seal 2, with the exception of the eyes and portions of the top of the head (see *Methods*). **(K)** Photographs of Seal 2 permit visual comparison between the real animal and CAD models for both head and whisker array. Panels A-F show data for N = 171 whiskers. Abbreviations: C: caudal; R: rostral; D: dorsal; V: ventral; CF: concave forwards; CB: concave backwards.

The subplots of Figs. 3DEF show that actual vs. predicted values for each angle of emergence are relatively uniformly dispersed about the equality line, confirming that the models of Figs. 3ABC were correctly chosen.

Visual intuition for whiskers’ orientation on the pad is provided in Figs. 3GHI. Most rostral whiskers emerge from the face pointing forwards and towards the midline, while most caudal whiskers emerge pointing slightly backwards (Fig. 3G). Dorsal whiskers emerge pointing upwards; ventral whiskers downwards (Fig. 3H). Whiskers in the dorsal-caudal regions of the array are oriented the most concave backwards, and rostral-ventral whiskers the most concave forwards (Fig. 3I). Finally, the pattern of variation for ζ_w_ is strikingly similar to that of θ_w_; in fact, these two variables are closely linear with r = 0.875 (Pearson correlation).

Together, Eqns 1 - 8 can be used to generate a full 3D model of the harbor seal vibrissal array. In Fig. 3J, three views of the 3D whisker array are superimposed on a 3D CAD model of the face and head of Seal 2. Comparing the model with the photographs of Seal 2 (Fig. 3K) provides visual intuition for the model’s descriptive ability. The equation based model with seal head is available at: WEBSITE.

### 3.3. Comparing vibrissal arrays of seals, mice, and rats

#### Whisker array size and shape

Equations for seals, rats, and mice are compared in Table 1 and visualized in Fig. 4. Interpreting these equations across species requires three comparisons. First, the equations’ functional forms indicate which independent variables 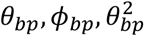 are statistically important. Second, the weight on each independent variable describes the variable’s role in the equation. A weight’s magnitude is unrelated to its statistical significance. Third, confidence intervals indicate whether weights are statistically the same across species, provided that the equations have the same form.

**Table 1.**
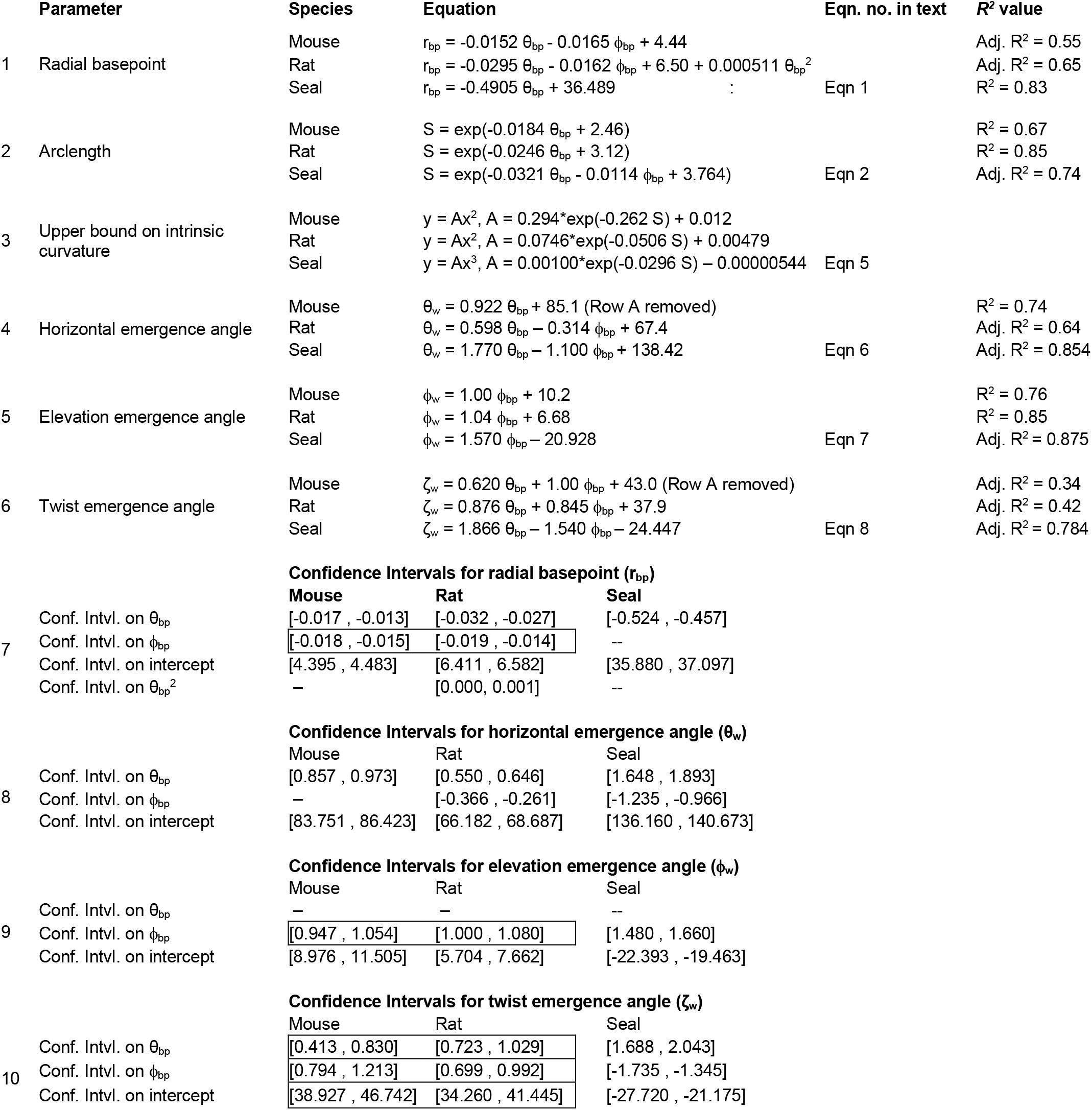
Equations for the geometry of the harbor seal vibrissal array are compared with those for rats and mice. Rat data from (Belli et al., 2018). Mouse data from (Bresee et al., 2023). All angles are in degrees and r_bp_ and S are in mm. The curvature coefficient (A) has units of 1/mm for mouse and rat, and 1/mm^2^ for the seal. Overlapping confidence intervals (Conf. Intvl.) are boxed.

**Fig. 4.**
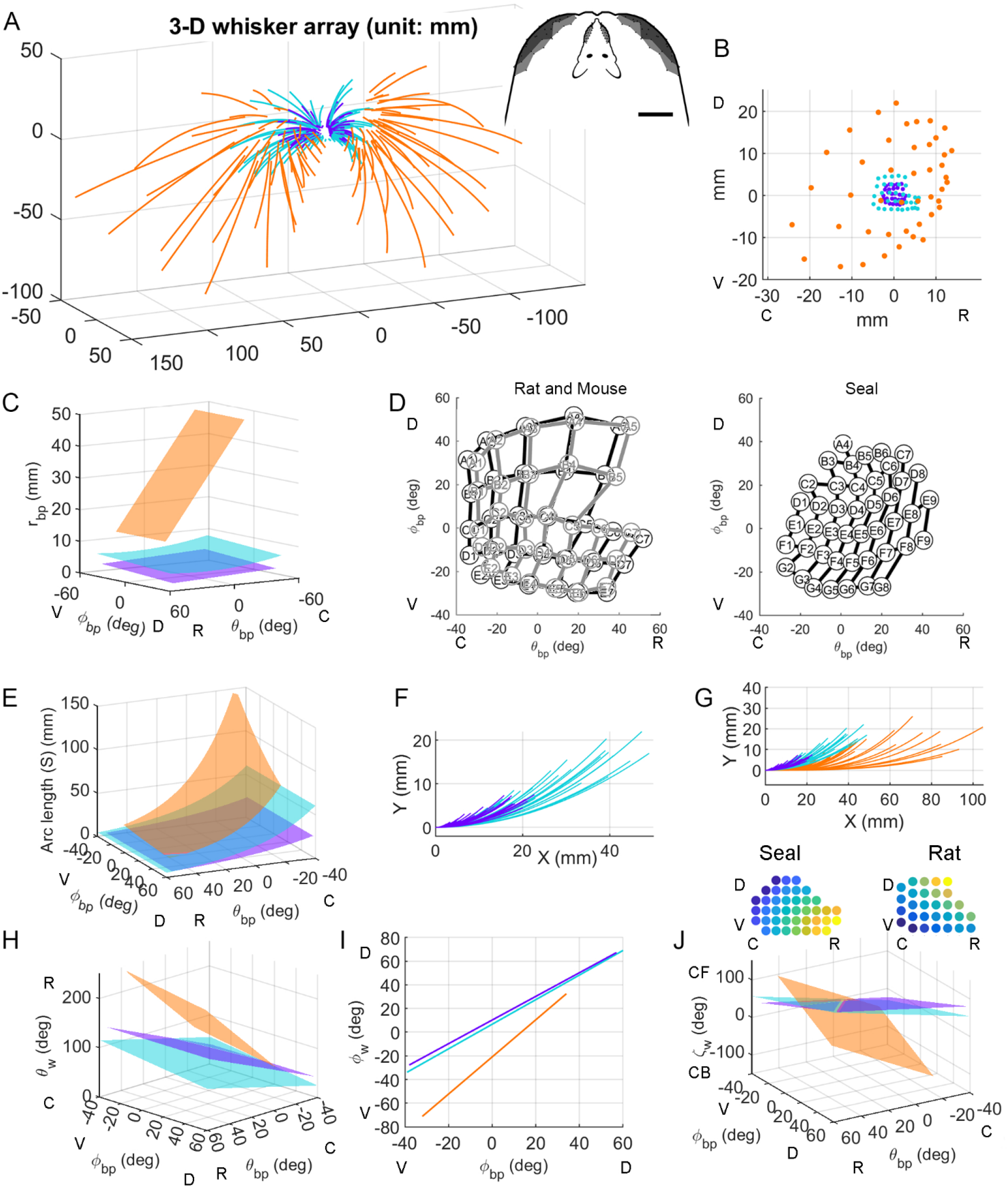
The harbor seal whisker array is compared to arrays of rats and mice. In all panels data for harbor seals are in orange, rats in teal, and mice in purple. Rat data are from (Belli et al., 2018). Mouse data are from (Bresee et al., 2023). **(A)** Full 3D geometric models of seal, rat, and mouse whisker arrays are shown in standard position and orientation. **(A, inset)** Top-down views compare rat and seal arrays. Scale bar is 20 mm. **(B)** Whisker basepoints are plotted in the sagittal (y-z) plane. **(C)** Plotting r_bp_ as a function of basepoint position highlights size differences between species. **(D)** To normalize for head size, mean whisker basepoint locations for all three species are plotted in θ_bp_ and ϕ_bp_. In these angular coordinates, seal whisker basepoints (right panel) are spaced more densely than mouse and rat whisker basepoints (overlayed in the left panel). The right panel shows the same data as Fig. 2B of the present work. **(E)** Whisker arclength (S) is plotted as a function of basepoint position (θ_bp_ and ϕ_bp_). **(F)** Seal whiskers are fit with cubic equations, while mouse and rat whiskers are fit with quadratic equations. **(G)** The large overlap in mouse and rat whisker shapes is revealed in an expanded view of (F). Proximal portions of all whiskers are aligned with the x-axis. **(H**,**I**,**J)** Rostral-caudal emergence angle (θ_w_), elevation emergence angle (ϕ_w_), and twist emergence angle (ζ_w_) are plotted as functions of basepoint position (θ_bp_ and ϕ_bp_). The top panels in (J) show heatmaps for the values of ζ_w_ in the bottom panel, for seal and rat. For both animals, the gradient of twist angles varies linearly with θ_bp_ and ϕ_bp_, but the gradient of the seal is orthogonal to the gradient of the rat. Abbreviations: C: caudal; R: rostral; D: dorsal; V: ventral; CF: concave forwards; CB: concave backwards.

The size and shape of the arrays of the three species are captured in Table 1 line 1, and in Fig. 4A-D. The form of the equation for r_bp_ is different between mouse, rat, and seal. For both mouse and rat, r_bp_ decreases as a function of θ_bp_ and ϕ_bp_, but the rat exhibits an additional dependence on 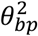, so confidence intervals (Table 1 line 7) cannot be directly compared. In the seal, r_bp_ decreases linearly with θ_bp_, with no ϕ_bp_ dependence, indicating that whisker distances from the origin form a surface approximately symmetric about the horizontal plane.

When 3D models of seal, rat, and mouse are overlaid (Fig. 4A) shape differences are less obvious than size differences. The entire rat whisker array fits within the bounds of the seal’s basepoints, and a top-down view (inset) reveals that nearly the entire rat head could fit between the seal’s most rostral left and right whiskers. A side-view of the basepoints (Fig. 4B) reinforces the large size difference. The seal’s basepoints encompass a surface area of ∼15 ±2.8 cm^2^, approximately 10 times larger than the area of the rat array(∼1.4 ±0.35 cm^2^; data from (Belli et al., 2018)), and 30 times larger than the mouse array (∼0.5 ±0.16 cm^2^; data from (Bresee et al., 2023)). Overall size and shape differences are also evident when r_bp_ is plotted as a function of θ_bp_ and ϕ_bp_ (Fig. 4C). This plots not only shows large size differences, but also that r_bp_ for the seal changes more steeply with θ_bp_, indicating that its whiskers are located more rostrally than those of the two rodents.

Although Figs. 4ABC show that seal whiskers are spaced further apart than rodent whiskers in linear distance, this effect could be due to head size. To normalize for head size, Fig. 4D plots basepoint spacing in angular coordinates, revealing that mice and rats have similar whisker densities, but the seal’s whiskers are far more densely spaced.

#### Arclength and intrinsic curvature

Consistent with results for mice and rats, seal whisker arclength (S) increases exponentially across the array (Table 1, line 2; Fig. 4E). However, unlike the two rodents, S depends on both rostrocaudal (θ_bp_) and dorsoventral (ϕ_bp_) whisker position. Because the weight for ϕ_bp_ is smaller than the weight for θ_bp_, the largest variation in S for the seal will occur in the rostrocaudal direction, as it does for rodents. The range of S is larger for rats than for mice, as seen in an overlay of all individual whisker fits (Fig 4F), and larger still for seals (Fig4EG). However, the longest seal whisker is only ∼2 times longer than the longest rat whisker, suggesting a restricted range of useful whisker lengths across body size.

The present work measured the intrinsic curvature of seal whiskers when they were hydrated, and uses single-parameter polynomial fits as intuitive descriptions of whisker shape. Rat and mouse whiskers are best fit by quadratics, while seal whiskers are best fit by cubics. These whisker shape differences are seen in the overlays of Fig 4FG, and could reflect adaptation to a hydrodynamic environment, differences in predation, or the large evolutionary separation between phocines and rodents. Like rat and mouse whiskers, longer seal whiskers tend to be straighter than shorter whiskers, and short whiskers exhibit large variability in curvature coefficient, with a sharp upper bound limiting the range (Table 1 line 3; Fig. 1G).

#### Emergence angles

Figs. 4HIJ visualize the equations for *θ*_*w*_, *ϕ*_*w*_, and *ζ*_*w*_ shown in lines 4, 5, and 6 of Table 1, with corresponding confidence intervals in lines 8, 9, and 10.

The values of the horizontal angle (*θθ*_*w*_) depend only on *θ*_*bp*_ for the mouse, but depend on *θ*_*bp*_ and *θθ*_*bp*_ for both rats and seals (Table 1, line 4). For all species, more rostral whiskers emerge at angles that point more rostrally, and for seals and rats, whiskers emerge more rostrally in ventral regions of the array. Values for *θ*_*w*_ are generally, but not always, larger in seals than in rodents (Fig. 4H), with the most rostral seal whiskers crossing 180° and pointing inward toward the midline.

A striking similarity between the whisker arrays of mice, rats, and seals is that the form of the equation for the whisker elevation angle (ϕ_w_) is identical for all three species (Table 1, line 5). There is a strong, single-variable, linear relationship between ϕ_w_ and the dorsoventral location of the whisker basepoint (ϕ_bp_). The relationship between ϕ_w_ and ϕ_bp_ for the seal has a significantly steeper slope and is offset towards negative values, indicating that whiskers emerge from the seal’s face pointing more ventrally than on mice or rats (Fig. 4I). The overlapping confidence intervals on the slope for mouse and rat (Table 1, line 9) indicate that the elevation angles of their whiskers change in the same way from dorsal to ventral.

A second important similarity is that the form of the equation for the whisker twist angle (ζ_w_) is identical for all three species (Table 1 line 6), exhibiting a linear dependence on both θ_bp_ and ϕ_bp_. All confidence intervals overlap for mouse and rat (Table 1, line 10), indicating that their twist angles are statistically indistinguishable.

Although the whisker twist angle (ζ_w_) varies diagonally across the array for all three animals, the direction of the diagonal is different, as clearly seen in the visualization of the equations for ζ_w_ (Fig. 4J). The surface for seal whiskers is oriented nearly orthogonally to the surfaces for rats and mice. The two figurines in the top panel plot the values of ζ_w_ as heat maps over the range of values in the bottom panel for the rat and seal. These heat maps clearly expose the near-orthogonal orientations of the gradients of twist angle for seal and rat.

## 4. Discussion

### 4.1. The head-centered approach permits important cross-species comparisons

The head-centered approach enables for the first time, comparisons of the 3D geometry of whisker arrays across species with different whisker numbers and arrangements. This approach serves as a foundation for cross-species geometric comparisons that may illuminate the functional use of whiskers during tactile sensing behaviors. The approach offers the first evidence for elements of a common whisker arrangement across species in different mammalian orders, suggesting conserved mechanisms of whisker-based tactile sensing. The approach also highlights differences across species, which may correlate with ethological specialization. Finally, the quantitative descriptions of array geometry inherent to the head-centered approach enable alignment with 3D CAD models for use in simulation and robotics experiments.

### 4.2. Cross-species similarities and differences in elevation and twist emergence angles

The whisker arrays of mice, rats, and seals share a linear relationship between elevation angle and the dorsoventral location of the whisker basepoint, indicating constant elevation within a row. This similarity is striking given that their last common ancestor likely lived between ∼90 and ∼65 MYA (Meredith et al., 2011; O’Leary et al., 2013). We suggest that this linear relationship may simplify calculating the elevation coordinate of a moving stimulus. When at rest, whiskers within a row will be contacted sequentially by a rostro-caudally moving stimulus. To combine information across whiskers and determine 3D stimulus location, the nervous system must track each whisker’s horizontal angle and twist angle separately, but only needs to track a single elevation angle per row. This advantage would also apply to mobile whiskers, assuming fixed relationships between protraction, elevation, and roll, as observed in rats and mice (Knutsen et al., 2008; Petersen et al., 2021).

A second similarity is the smooth diagonal gradient of whisker twist across the array, which may aid in determining direction of stimulus motion. Stimuli at different pitches relative to the animal’s head will encounter different gradients of whisker twist. Stimulus direction could be inferred from these gradients because the mechanical response of each whisker will depend strongly on its orientation relative to the stimulus (Quist and Hartmann, 2012; Yu et al., 2016b).

Seal 3D geometry also differs from that of the two rodents. In seals, ventral-rostral whiskers tend to be oriented concave downwards; this orientation shifts diagonally across the array to become concave backwards in dorsal-caudal regions. In rats and mice, dorsal-rostral whiskers are concave forwards, transitioning diagonally to concave downwards in ventral-caudal regions. These orientations make sense ethologically – pushing concave-forwards whiskers through water would cause turbulence, complicating sensing. In contrast, rotating concave-forward whiskers through air extends tactile reach. A second difference between species is that only seals exhibit a linear relationship between θ_w_ and ζ_w_, discussed further below.

### 4.3. Quantifying 3D whisker arrangement for the harbor seal: mechanical consequences for tracking wakes

Harbor seals use whiskers to track fish wakes (Dehnhardt et al., 1998; Dehnhardt et al., 2001; Wieskotten et al., 2010; Wieskotten et al., 2011). Several findings of the present work have direct mechanical consequences for wake tracking.

Basic physical principles dictate that the regular variation in arclength across the array (Fig. 2D-F) will create a mechanical filter (Hanke et al., 2010), since the natural frequencies of a tapered cantilever beam depend on the ratio of its base diameter to its arclength squared (Georgian, 1965). Shorter (longer) whiskers will tend to respond more strongly to higher (lower) frequency components of vortices, wakes, and direct tactile stimuli. At the same time, undulations along the whisker length will tend to reduce vortex-induced vibrations of the whisker, thereby improving tracking (Ginter et al., 2012; Hanke et al., 2010).

The present work showed that the orientation of the intrinsic curvature of whiskers varies diagonally across the array (Fig. 3CFI). This “curvature orientation” gradient is likely already sufficient to ensure that stimuli at different pitches relative to the animal’s head will generate different mechanical gradients. However, the elliptical cross-sections of seal whiskers may enhance this effect, as a whisker’s vibrations will be larger when the wide, rather than thin, diameter of the whisker faces into the flow (Murphy et al., 2013). Although the present work did not measure whisker diameters, (Summarell et al., 2015), found that the intrinsic curvature of a harbor seal whisker aligns with its major axis and that flexural stiffness along the major axis is nearly double that of the minor axis. The orientation of greatest stiffness will thus correlate with the twist angles shown in Figs. 3CFI, likely increasing response magnitude in the direction of the gradient and reducing disturbances orthogonal to it. Because “curvature orientation” and “diameter orientation” are correlated, future work will need to avoid confounding their mechanical effects.

For a seal swimming forwards, changes in θ_w_ change the angle of attack of the whisker, effectively changing the area projected into the flow, and thus how the whisker bends and vibrates. Intriguingly, for the seal – but not the rat – the two angles θ_w_ and ζ_w_ are strongly correlated (Figs. 3GI). By varying these two angles virtually in lock step across the array, the seal vibrissal system may ensure that these two different sources of vibration variability are always correlated. Rats and mice may not require such a correlation, which could explain why the ζ_w_ gradient varies in different directions between the seal and the two rodents.

### 4.4. Models of animal biomechanics in neuroscience and robotics

The mechanics of wake-following behavior strongly depend on the seal’s active tracking motions, which likely change based on the external wakes it senses. Modeling this closed-loop sensorimotor behavior, and the neural circuitry that underlies it, will require the types of 3D quantification and CAD models presented here. This work will advance the study of the biomechanics and neurobiology of wake detection and tracking, as well as the development of underwater seal-like robots. In fact, the seal array model developed here has already been used to explore how whisker signals reflect flow structure (Liu et al., 2023).

## ACKNOWLEDGMENTS

We thank Dr. Christine Scholtyßeck for help handling the seals. E. Claire Mosier, an undergraduate student in Northwestern’s Biomedical Engineering department, helped us take some of the data in Rostock. We are grateful to RAPIDFORM™ for providing a temporary stand-alone license to acquire and analyze the scans when we briefly lost access to the internet at a critical point in the work. We thank Yifu Luo for remaking several of the figures. Portions of this work have been published in two Ph.D. dissertations: Graff, MM (2016) *Mechanisms of Whisker Based Flow Sensing*, Ph.D. Thesis, Northwestern University. ProQuest/UMI, 130 pages (Publication No. 10160640). Belli, HM (2017) *Sensing with Whiskers: From Geometry and Mechanics to the Statistics of the Array*, Ph.D. Thesis, Northwestern University, ProQuest/UMI, 253 pages (Publication No. 10681095).

