## Supplementary Table 1 for "Spatial arrangement of the whiskers of harbor seals (*Phoca vitulina*) compared to whisker arrangements of house mice (*Mus musculus*) and brown rats (*Rattus norvegicus*)"

**Supplementary Table S1.** Best fit equations for radial basepoint coordinate and angles of emergence by individual animal, along with combined equations across animals. *Abbreviations:* Prm: Parameter; (n): number of whiskers.

| | Prm | Animal (n) | Coefficients $c_1$ | $c_2$ | $c_3$ | 95% Confidence Intervals $c_1$ | $c_2$ | $c_3$ | Adj. $R^2$ |
|---|---|---|---|---|---|---|---|---|---|
| 1 | $r_{bp}$ | Seal 1 (56) | N/A | -0.42203 | 36.525 | N/A | [-0.4729, -0.3711] | [35.5827, 37.4682] | 0.833 |
| | | Seal 2 (62) | 0.003283 | -0.53157 | 35.899 | [4.970e-04, 0.0060702] | [-0.5945, -0.4687] | [34.6273, 37.1707] | 0.838 |
| | | Seal 3 (53) | 0.004162 | -0.61645 | 35.195 | [0.001343, 0.006980] | [-0.6877, -0.5452] | [33.8951, 36.4957] | 0.882 |
| | | Combined (171) | N/A | -0.4905 | 36.489 | N/A | [-0.5236, -0.4574] | [35.8804, 37.0967] | 0.846 |

Final equation for radial basepoint coordinate: $r_{bp} = c_1\theta_{bp}^2 + c_2\theta_{bp} + c_3$, where $r_{bp}$ is in millimeters and $\theta_{bp}$ is in degrees.

| | Prm | Animal (n) | $c_1$ | $c_2$ | $c_3$ | $c_4$ | $c_1$ | $c_2$ | $c_3$ | $c_4$ | Adj. $R^2$ |
|---|---|---|---|---|---|---|---|---|---|---|---|
| 2 | $\theta_w$ | Seal 1 (56) | N/A | -0.972 | 1.815 | 133.02 | N/A | [-1.1597, -0.7849] | [1.6454, 1.9849] | [129.9012, 136.1351] | 0.903 |
| | | Seal 2 (62) | N/A | -1.141 | 1.647 | 138.42 | N/A | [-1.3181, -0.9631] | [1.4731, 1.8199] | [135.3208, 141.5222] | 0.889 |
| | | Seal 3 (53) | 0.0504 | -0.749 | 1.856 | 133.85 | [0.0394, 0.0615] | [-0.9656, -0.5319] | [1.6882, 2.0236] | [129.8400, 137.8607] | 0.923 |
| | | Combined (171) | N/A | -1.100 | 1.770 | 138.42 | N/A | [-1.2346, -0.9655] | [1.6481, 1.8927] | [136.1603, 140.6733] | 0.854 |

Final equation for azimuthal angle of emergence: $\theta_w = c_1\phi_{bp}^2 + c_2\phi_{bp} + c_3\theta_{bp} + c_4$, where $\theta_w$, $\phi_{bp}$ and $\theta_{bp}$ are in degrees.

| | Prm | Animal (n) | $c_1$ | $c_2$ | $c_3$ | $c_4$ | $c_1$ | $c_2$ | $c_3$ | $c_4$ | Adj. $R^2$ |
|---|---|---|---|---|---|---|---|---|---|---|---|
| 3 | $\phi_w$ | Seal 1 (56) | N/A | 1.700 | N/A | -23.345 | N/A | [1.5198, 1.8800] | N/A | [-26.3311, -20.3596] | 0.867 |
| | | Seal 2 (62) | N/A | 1.449 | N/A | -21.738 | N/A | [1.3460, 1.5526] | N/A | [-23.4949, -19.9809] | 0.928 |
| | | Seal 3 (53) | 0.0127 | 1.730 | N/A | -19.799 | [0.003515, 0.02179] | [1.5494, 1.9098] | N/A | [-22.9582, -16.6397] | 0.882 |
| | | Combined (171) | N/A | 1.570 | N/A | -20.928 | N/A | [1.4802, 1.6597] | N/A | [-22.3935, -19.4632] | 0.875 |

Final equation for elevation angle of emergence: $\phi_w = c_1\phi_{bp}^2 + c_2\phi_{bp} + c_3\theta_{bp} + c_4$, where $\phi_w$, $\phi_{bp}$, and $\theta_{bp}$ are in degrees.

| | Prm | Animal (n) | $c_1$ | $c_2$ | $c_3$ | $c_4$ | $c_1$ | $c_2$ | $c_3$ | $c_4$ | Adj. $R^2$ |
|---|---|---|---|---|---|---|---|---|---|---|---|
| 4 | $\zeta_w$ | Seal 1 (56) | N/A | -1.350 | 1.573 | -23.663 | N/A | [-1.7195, -0.9803] | [1.2382, 1.9078] | [-29.8110, -17.5155] | 0.693 |
| | | Seal 2 (62) | N/A | -1.565 | 1.753 | -23.714 | N/A | [-1.8397, -1.2893] | [1.4836, 2.0214] | [-28.5220, -18.9053] | 0.818 |
| | | Seal 3 (53) | N/A | -1.732 | 2.308 | -28.091 | N/A | [-2.1166, -1.3482] | [1.9916, 2.6238] | [-34.2624, -21.9194] | 0.840 |
| | | Combined (171) | N/A | -1.540 | 1.866 | -24.447 | N/A | [-1.7351, -1.3448] | [1.6882, 2.0429] | [-27.7197, -21.1746] | 0.784 |

Final equation for twist angle of emergence: $\zeta_w = c_1\phi_{bp}^2 + c_2\phi_{bp} + c_3\theta_{bp} + c_4$, where $\zeta_w$, $\phi_{bp}$, and $\theta_{bp}$ are in degrees.
